# Derivation and Characterization of an inositol phosphate-independent HIV-1

**DOI:** 10.1101/2021.03.01.433343

**Authors:** Daniel Poston, Trinity Zang, Paul Bieniasz

## Abstract

A critical step in the HIV-1 replication cycle is the assembly of Gag proteins to form virions at the plasma membrane. Virion assembly and maturation is facilitated by the cellular polyanion inositol hexaphosphate (IP_6_), which is proposed to stabilize both the immature Gag lattice and the mature capsid lattice by binding to rings of primary amines at the center of Gag or capsid protein (CA) hexamers. The amino acids comprising these rings are critical for proper virion formation and their substitution results in assembly deficits or impaired infectiousness. To better understand the nature of the deficits that accompany IP_6_-deficiency, we passaged HIV-1 mutants that had substitutions in IP_6_-coordinating residues to select for compensatory mutations. We found a mutation, a threonine to isoleucine substitution at position 371 (T371I) in Gag, that restored replication competence to an IP_6_-binding-deficient HIV-1 mutant. Notably, unlike wild-type HIV-1, the assembly and infectiousness of resulting virus was not impaired when IP6 biosynthetic enzymes were genetically ablated. Surprisingly, we also found that the maturation inhibitor Bevirimat (BVM) could restore the assembly and replication of an IP_6_-binding deficient mutant. Moreover, using BVM-dependent mutants we were able to image the BVM-inducible assembly of individual HIV-1 particles assembly in living cells. Overall these results suggest that IP_6_-Gag and Gag-Gag contacts are finely tuned to generate a Gag lattice of optimal stability, and that under certain conditions BVM can functionally replace IP_6_.

**Author Summary:** A key step in HIV-1 replication is the assembly of virions that are released from the infected cell. Previous work has suggested that a small molecule called IP_6_ is critical role in this process, promoting both HIV-1 assembly and the stability of mature fully infectious virions. Since IP_6_ is required for multiple steps in HIV-1 assembly and maturation, it is a candidate for the development of anti-retroviral therapies. Here, we identify an HIV-1 mutant that replicates independently of IP_6_, and show that a different small molecule can functionally substitute for IP_6_ under certain conditions. These findings suggest that IP_6_ regulates the stability of protein interactions during virion assembly and that the precise degree stability of these interactions is finely tuned and important for generating infectious virions. Finally, our work identifies an inducible virion assembly system that can be utilized to visualize HIV-1 assembly events using live cell microscopy.

## Introduction

The HIV-1 Gag polyprotein which is composed of the matrix (MA), capsid (CA), spacer Peptide 1 (SP1), nucleocapsid (NC), spacer Peptide 2 (SP2), and p6 domains, has central critical structural and functional roles in the HIV-1 replication cycle. During virion assembly, multimerization of the Gag polyprotein at the plasma membrane, primarily driven by the CA and NC domains, generates immature HIV-1 virions composed of radially oriented Gag hexamers (1–5). Following assembly, and concomitant with or shortly after nascent particles are released, proteolytic processing of Gag by HIV-1 protease separates the aforementioned Gag domains (6). The liberated CA protein undergoes a major structural rearrangement to form the mature conical core, composed of a lattice of CA hexamers with 12 CA pentamers, and is a salient feature of maturation (7). Only after maturation are HIV-1 particles able to initiate new cycles of infection.

It has been previously shown that inositol phosphates play a critical role in both HIV-1 assembly and maturation. While assembly of HIV-1 Gag protein *in vitro* yields immature particles that differ in size and character from authentic virions, addition of inositol phosphates to *in vitro* assembly reactions enables the production of particles that resemble authentic virions (8). Further work identified inositol hexakisphosphate, or IP_6_, as the key mediator of this process. IP_6_, is a ubiquitous cellular polyanion containing 5 equatorial phosphates and a single axial phosphate, and facilitates formation of immature HIV-1 Gag lattice by binding to and stabilizing positively-charged rings of primary amines. These rings are formed by lysine residues at Gag positions 290 and 359 (K290 & K359) that are positioned at the center of the immature Gag hexamer (9). Following the subsequent structural rearrangement of CA that accompanies maturation, IP_6_ next binds to a second, distinct positively charged ring in the mature CA hexamer formed by arginine residues at CA position 18 (R18, Gag position R150). The R18 ring stabilizes the mature CA hexamer, and is required for viral DNA synthesis in newly infected cells (10–12). It is thought that IP_6_ is recruited into virions by interacting with K290 and K359 during immature particle production; this model is consistent with data demonstrating that HIV-1_K290A_ and HIV-1_K359A_ are significantly impaired in both viral production and IP_6_ packaging, while HIV-1_K359I_ is assembly competent but generates poorly infectious particles (13).

The importance of each of the IP_6_-coordinating residues has been established, as mutagenesis of any such residue to an alanine (HIV-1_R18A,_ HIV-1_K290A_, or HIV-1_K359A_) significantly impairs infectivity in either single cycle or spreading infection assays (9,13). Additionally, yield of infectious virions is also substantially reduced in cells lacking key enzymes in the IP_6_ biosynthetic pathway (IPPK or IPMK) or in cells overexpressing MINPP1, a phosphatase that dephosphorylates IP_6_ (9,13,14). The IP_6_-coordinating amino acids are conserved among diverse lentiviruses, suggesting a general requirement for IP_6_ (15).

While there is considerable evidence that perturbing IP_6_ binding impairs HIV-1 replication, further investigation into the precise mechanisms underlying replication deficits is warranted. To better understand the role of IP_6_, we serially passaged virions containing substitutions in IP_6_-coordinating residues to identify second-site compensatory mutations that might rescue the resulting infectivity deficits. Accordingly, we found a single substitution that rescued the replication deficit observed in two IP_6_ binding-deficient mutants. Using CRISPR/Cas9 knockout of IPMK, we show that the second-site substitution renders infectious virion yield independent of cellular IP_6_ levels. Strikingly, we also found that treatment with a maturation inhibitor Bevirimat (BVM) rescues infectivity of the IP_6_-binding-deficient mutant HIV-1_K359A_, indicating that BVM can functionally substitute for IP_6_ in certain situations. Indeed, using approaches in which the assembly of individual HIV-1 particles is imaged in living cells, we show that addition of BVM can induce the assembly of CA-mutant HIV-1 virions.

## Results

### Identification of a second-site substitution that restores replication competence to IP_6_-binding deficient HIV-1 mutants

To identify second site changes that would rescuing IP_6_ deficiency, we passaged HIV-1 mutants encoding substitutions in IP_6_ coordinating residues (HIV-1_R18A_, HIV-1_K290A,_ or HIV-1_K359A_) in the highly-permissive MT4 T-cell line. Initial attempts in which MT4 cells were infected with mutant viral stocks were unsuccessful, likely due to the dramatically impaired fitness of these mutants and consequent inability to establish a sufficiently large population of infected cells to generate revertants. To overcome this problem, we instead co-cultured MT4 cells with virus-producing 293T cells that had been transfected with HIV-1_R18A_, HIV-1_K290A,_ and HIV-1_K359A_ proviral plasmids that encode GFP in place of nef. After removing the 293T cells, infected MT4 cells were co-cultured with uninfected MT4 cells, until most of the MT4 cells became infected (as monitored by visual inspection of GFP+ cells in the culture). Thereafter, cell-free supernatant was serially passed in MT4 cells (Figure 1A).

**Fig 1.**
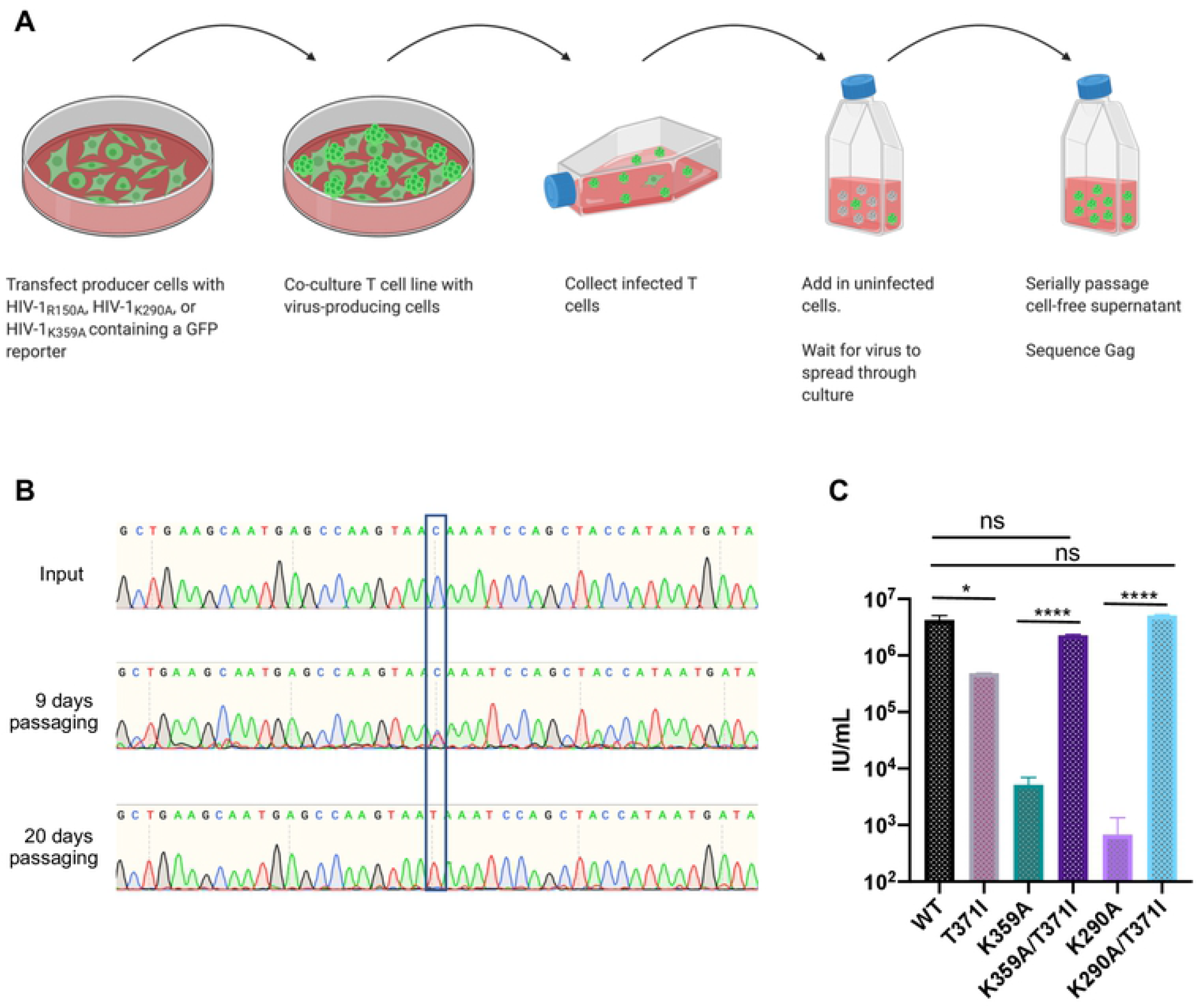
Derivation of an IP6-independent HIV-1 mutant. (A) Passaging schematic. (B) Sanger sequencing of HIV-1 gag indicating emergence of a single-site revertant. Input or viral RNA was isolated at indicated timepoint, and the gag region was amplified by RT-PCR. (C) Single cycle infection assay confirming the identified substitution rescues infectivity of K359A. 293Ts were transfected with proviral clones of WT HIV-1_NHG_ or indicated mutant, and 48 hours post transfection, supernatant was titrated on MT4 cells and assessed using flow cytometry. Statistical analysis: Student’s T test.

For one mutant, HIV-1_K359A_, observation of GFP positive cells suggested that a apparently compensatory mutation arose after approximately 2 weeks of passaging. PCR amplification and sequencing of Gag encoding sequences from this culture revealed a single nucleotide substitution that resulted in a threonine to isoleucine substitution at Gag position 371 (Figure 1B). No revertant mutants could be obtained for HIV-1_R18A_ or HIV-1_K290A_. This finding may reflect a greater magnitude of impairment of these particular substitutions, making the generation of revertant mutants more difficult.

To determine whether the T371I mutant rescued the infectivity defect present in HIV-1_K359A_, we generated a proviral clone, HIV-1_K359A/T371I_, encoding both mutations and measured the infectious virion yield from proviral plasmid-transfected 293T cells. Addition of the T371I substitution to HIV-1_K359A_ restored infectious virion yield to wild-type levels (Figure 1C). Although this second-site, apparently compensatory change was identified only in the context of HIV-1_K359A_, we asked whether the T371I substitution could rescue the HIV-1_K290A_, given purported similar roles of K290 and K359 in binding IP_6_. Indeed, found that HIV-1_K290A/T371I_, unlike HIV-1_K290A_, yielded similar levels of infectious HIV-1 virions to wild-type HIV-1.

### Infectious HIV-1_K359A/T371I_ particle yield is not affected ablation of IP_6_ synthesis in virus producing cells

Because the HIV-1_K359A_ is defective for IP_6_ binding we next asked whether HIV-1_K359A/T371I_ retained infectiousness when cellular IP_6_ levels were reduced. Using CRISPR/Cas9 we generated 293T cell lines lacking IPMK, an enzyme in the IP_6_ synthetic pathway. Previous work has demonstrated IPMK knockout cells have greatly reduced levels of both IP_5_ and IP_6_ (13). To account for potential clonal variation in capacity to generate HIV-1 particles, we used 3 separate IPMK targeting sgRNAs or a corresponding empty vector to generate ten independent single cell clones of IPMK knockout and WT control 293T cells (Figure 2A). The loss of IPMK was confirmed by DNA sequencing of target loci, which revealed the introduction of frameshift mutations into both alleles of the IPMK coding sequences, and the absence of intact IPMK alleles. In agreement with previous studies, the yield of infectious HIV-1_WT_ virions from IPMK-deficient 293T cells was significantly decreased, by 10-fold (p=0.0091, Figure 2A). The yield of HIV-1_K359A_ from 293T cells was greatly reduced compared to wildtype HIV-1 as expected, and was not further reduced by IPMK deficiency (Figure 2B). Importantly, the yield of HIV-1_K359A/T371I_ was only marginally reduced compared to wild type HIV-1 and there was no difference in yield of infectious HIV-1_K359A/T371I_ from WT 293T cells versus IPMK deficient 293T cells (Figure 2C, p=0.178).

**Fig 2.**
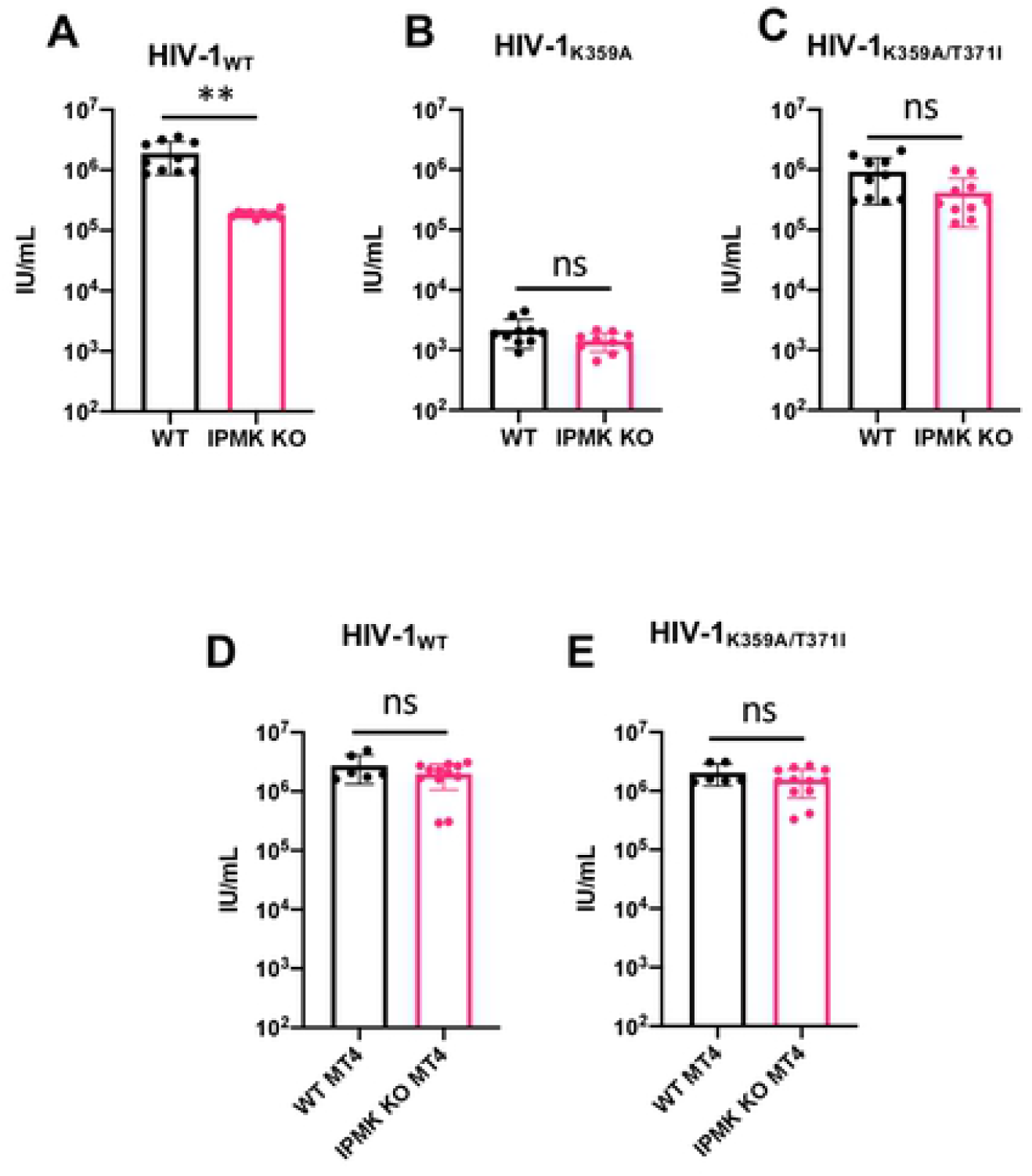
HIV-1_K359A/T371I_ is not impaired by lack of cellular IP6. (A-C): Control or IPMK KO single 293T cell clones were transfected with HIV-1_WT_ (A), HIV-1_K359A_ (B), or HIV-1_K359A/T371I_ (C) proviral plasmids. After 48 hours, supernatants were collected and titrated on MT4 cells. Each data point represent a different 293T cell clone. Statistical analysis: unpaired Student’s t-test. (D-E) HIV-1_WT_ (D) or HIV-1_K359A/T371I_ (E) virions were titrated on WT MT4 or IPMK KO MT4 cells and infection quantified by flow cytometry. Each data point represent a different MT4 cell clone. Statistical analysis: unpaired Student’s t-test.

### Neither HIV-1_WT_ nor HIV-1_K359A/T371I_ require IP_6_ synthesis in target cells during single cycle infection

It has been proposed that residues K290 and K359 selectively recruit IP_6_ into HIV-1 virions during assembly, thereby providing the source of the IP_6_ that binds to and stabilizes the R18 ring in the mature capsid core. The rationale for this idea stems from previous studies which have demonstrated that reduction of cellular IP_6_ levels in target cells does not impact susceptibility to incoming infection (13). Because HIV-1_K359A/T371I_ is fully infectious despite encoding a mutation that diminishes IP_6_ packaging into virions, we next asked whether HIV-1_K359A/T371I_ requires IP_6_ in target cells to be maximally infectious. We generated multiple twelve IPMK-deficient MT4 target cell clones and six control clones and performed single cycle infection assays using HIV-1_WT_ and HIV-1_K359A/T371I_ (Figure 2D-E). In agreement with previous studies (13), there was no difference in the infectiousness of HIV-1_WT_ in WT or IPMK-deficient MT4 cells (Figure 2D p = 0.3863). Moreover, there was no deficit in the infectiousness of HIV-1_K359A/T371I_ in WT or IPMK-deficient MT4 target cells (Figure 2E, p=0.4331).

### Bevirimat rescues infectious virion formation by the IP_6_-binding deficient mutant HIV-1_K359A_

Notably, The T371I mutation identified herein had been described previously in a different context. Specifically, this substitution was reported to stabilize the immature CA-SP1 lattice, mimicking the effect of maturation inhibitors (MI) (16,17). Therefore, we next asked whether maturation inhibitors themselves could rescue the deficit in infectious virion yield exhibited by HIV-1_K359A_. As a control, we included the previously described assembly-defective, maturation inhibitor-dependent CA mutant HIV-1_P289S_. We found that that BVM indeed rescued the infectivity of HIV-1_K359A_and HIV-1_P289S_ in both single-cycle and spreading replication assays. Specifically, in single cycle assays, BVM increased the yield of infectious HIV-1_K359A_ virions, up to 50-fold, and in a dose-dependent manner (Figure 3A). In spreading replication assays, BVM restored HIV-1_K359A_ spreading replication to levels similar to that of BVM-treated wildtype virus (Figure 3B).

**Fig 3.**
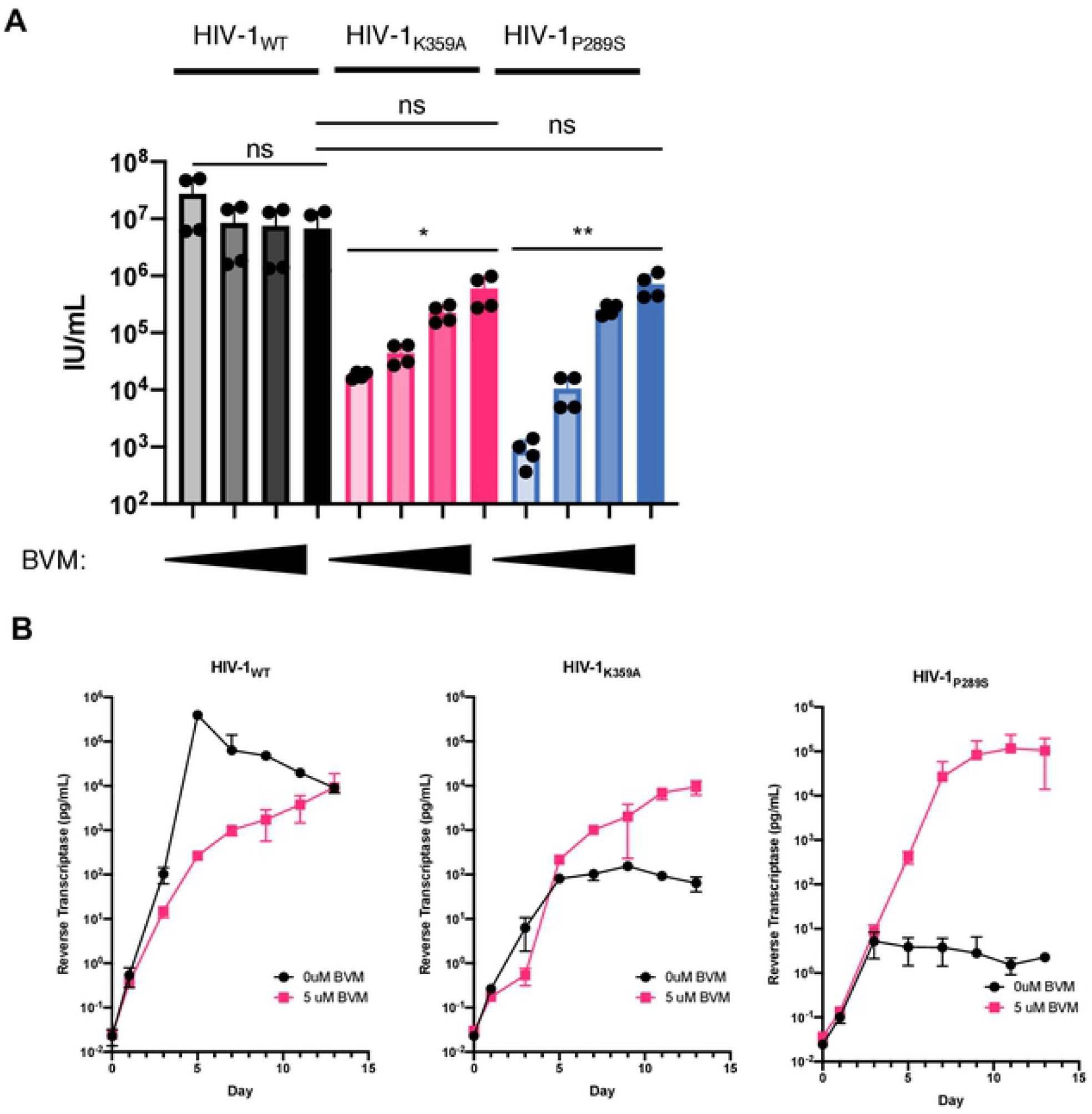
Bevirimat rescues infectivity of HIV-1_K359A_. (A) 293T cells were transfected with HIV-1_WT_ or HIV-1_K359A_ proviral plasmids in the presence of 0, 1, 5, or 10 μM Bevirimat. After 48 hours, supernatants were collected and titrated on MT4 cells and infection quatified by flow cytometry. (B) MT4 cells were infected with HIV-1_WT_ or HIV-1_K359A/T371I_ at an MOI of .01. At 16 hours post infection, inoculum was washed away and supernatants were collected every 48 hours for 13 days. Reverse transcriptase activity was measured in the supernatant samples using SYBR-PERT.

### BVM increases release of HIV-1_K359A_ virions independently of the viral protease

The interaction between K359 amines and IP_6_ likely stabilizes the immature Gag lattice, Similarly, maturation inhibitors are known bind to the immature Gag hexamers at approximal site and stabilize the immature CA-SP1 lattice (18,19). Therefore, we hypothesized that BVM rescue particle formation by HIV-1_K359A_ by stabilizing an otherwise destabilized lattice, effectively serving as a functional replacement for IP_6_. To test this idea, we measured the release of HIV-1_K359A_ virions from BVM-treated 293T cells by western blotting. BVM indeed increased the yield of HIV-1_K359A_ virions, in a dose-dependent manner (Figure 4A). To confirm that this effect was not due to BVM-mediated inhibition of Gag proteolysis, we performed similar experiments in virions containing an inactivating mutation in protease. We observed a similar dose-dependent increase in immature particle release, even in the context of protease inactivation (Figure 4B), suggesting that the effect of BVM on HIV-1_K359A_ assembly is due to the direct effect of BVM on the immature lattice, not through inhibition of proteolytic cleavage at the Gag-SP1 junction.

**Fig 4.**
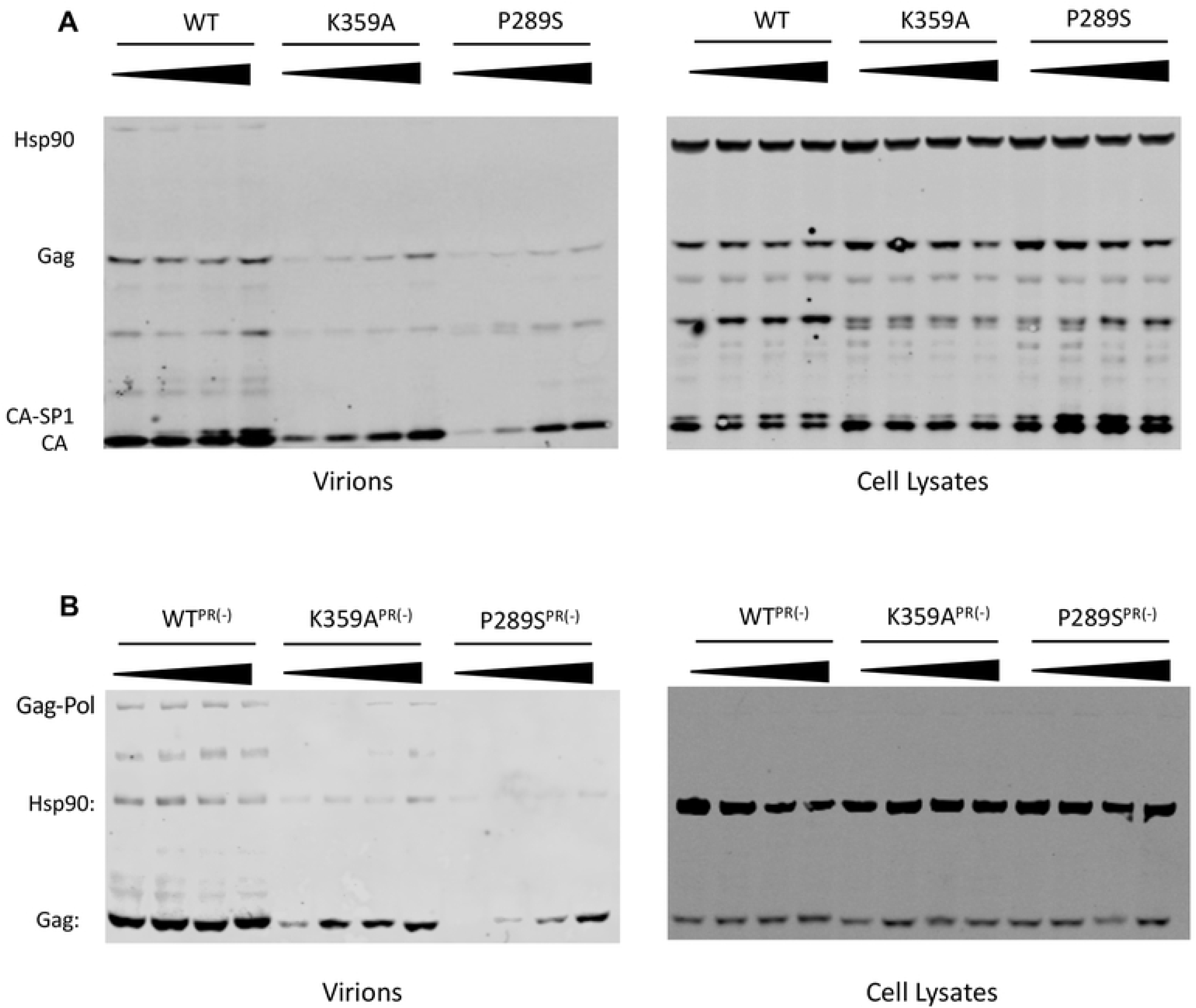
Bevirimat increase release of HIV-1_K359A_ independently of protease inhibition. (A) 293T cells were transfected with HIV-1_WT_, HIV-1_K359A,_ or HIV-1_P289S_ in the presence of 0, 1, 5, or 10 μM Bevirimat. 48 hours post transfection, supernatan (B) 293T cells were transfected with HIV-1_WT_, HIV-1_K359A,_ or HIV-1_P289S_ bearing an inactivating mutation in protease in the presence of 0, 1, 5, or 10 μM Bevirimat and analyzed via SDS-PAGE as above.

### Visualization of BVM-induced HIV-1 assembly observed in real time using live cell fluorescence microscopy

The above data strongly suggested that BVM rescues infectivity and release of HIV-1_K359A_ by facilitating particle assembly. To directly observe effects on virion assembly, we performed fluorescence microscopy using a novel imaging construct based on HIV-1_NL4-3_, in which Pol has been replaced by an HIV-1 codon-mimicking mNeonGreen, and in which Env and Vpu bear inactivating mutations. The resulting construct, herein referred to as HIV-1 NG, generates Gag-mNeonGreen in place of Gag-Pol during a single cycle of infection, thus allowing visualization of particle assembly as punctae at the plasma membrane.

We first infected TZM-bl cells with this reporter and derivatives (HIV-1 NG_WT_, HIV-1 NG_K359A_, and HIV-1 NG_P289S_) in the absence or presence of 5 μM BVM and performed widefield imaging on fixed cells 48 hours post infection. As indicated in Figure 5A, in the absence of BVM, cells infected with HIV-1 NG_K359A_ and HIV-1 NG_P289S_ exhibited primarily diffuse cytoplasmic fluorescence, and fewer punctae than for HIV-1 NG_WT_ infected cells, indicating impaired assembly. However, when infections were done in the presence of BVM, there were clearly increased numbers of membrane associated punctae in HIV-1 NG_K359A_ and HIV-1 NG_P289S_ infected cells suggesting BVM is able to directly facilitate particle assembly by these mutants.

**Fig 5.**
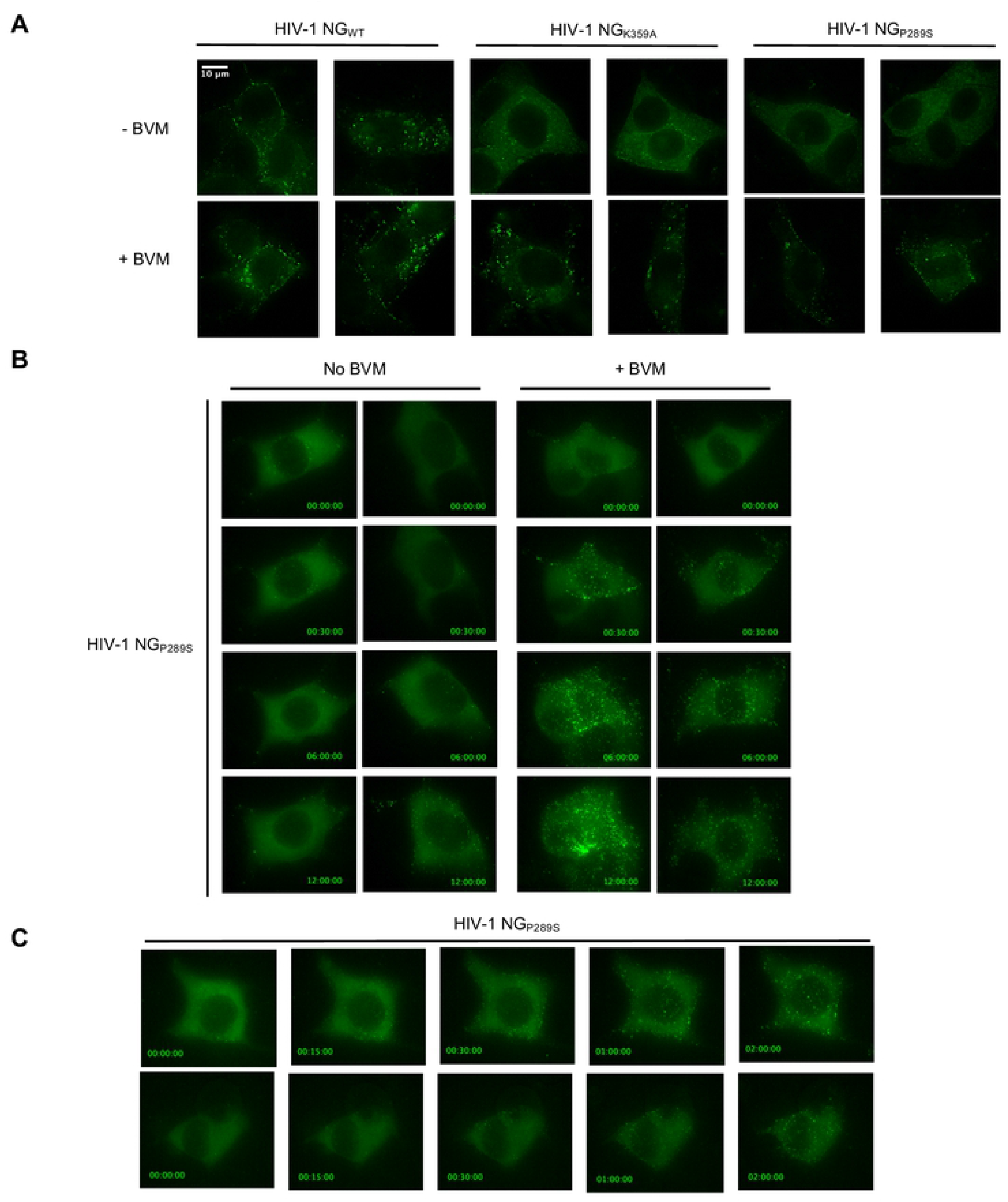
Visualization of Bevirimat-Induced assembly by fluorescence microscopy. (A) Fluorescence microscopy of TZM cells infected with a reporter HIV-1 virus encoding mNeon Green in place of Pol (HIV-1 NG) to visualize assembly. Representative images 48 hours-post-infection of HIV-1 NG_WT_, HIV-1 NG_K359A_, or HIV-1 NG_P289S_ infected in the absence or presence of 5μM BVM. (B) Representative time lapse images of TZM cells infected with HIV-1 NG_P289S_ -/+ treatment with 5μM BVM at 26 hours post infection. Image acquisition began immediately after BVM addition. Image labels: Hours:Minutes:Seconds post BVM addition (C) Representative time-lapse images of TZM cells treated with BVM at 26 hours post infection with HIV-1 NG_P289S._ Image labels: Hours:Minutes:Seconds post BVM addition.

This ability to induce HIV-1 particle assembly via addition of an exogenous small molecule has potential applications in imaging and other studies, as an inducible particle assembly system. In order to test the possible utility of this approach, we performed live cell widefield imaging studies using HIV-1 NG_P289S_, as this mutant displayed a greater responsiveness to BVM-induced assembly (see Figures 3A, 3B, and 4B). We infected TZM-bl cells in the absence of BVM and then, at 26 hours after infection, added BVM 5 μM and began acquiring images at 30 minute intervals. In the absence of BVM, few punctae are apparent in HIV-1 NG_P289S_ infected cells, even after 12 hours of imaging (Figure 5B, S1 Movie, S2 Movie). However, in the BVM-treated cells, substantially more punctae were evident, as soon as 30 minutes after BVM addition (Figure 5B, S3 Movie, S4 Movie).

Given that such striking differences could be observed as early as 30 minutes post BVM addition, we repeated these experiments with increased time resolution. TZM-bl cells were infected with HIV-1 NG_P289S_ for 26 hours and treated with BVM as above, followed by immediate image acquisition at 3 minute intervals. Assembly of HIV-1_P289S_ was rapidly induced by BVM (Figure 5C, S5 Movie, S6 Movie) with substantial numbers of punctae forming within 30-120min. We performed similar experiments using TIRF-FM imaging and quantified the presence of Gag-NG punctae over time, with similar results. Specifically, in the absence of BVM there were very few punctae evident at the plasma membrane (Figure 6, S7 Movie, 8 Movie). However, shortly after the addition of BVM, numerous punctae rapidly formed at the plasma membrane (Figure 6, S9 Movie, S10 Movie). These data provide proof of principle that such a system could be used to experimentally manipulate HIV-1 assembly for imaging or functional studies.

**Fig 6.**
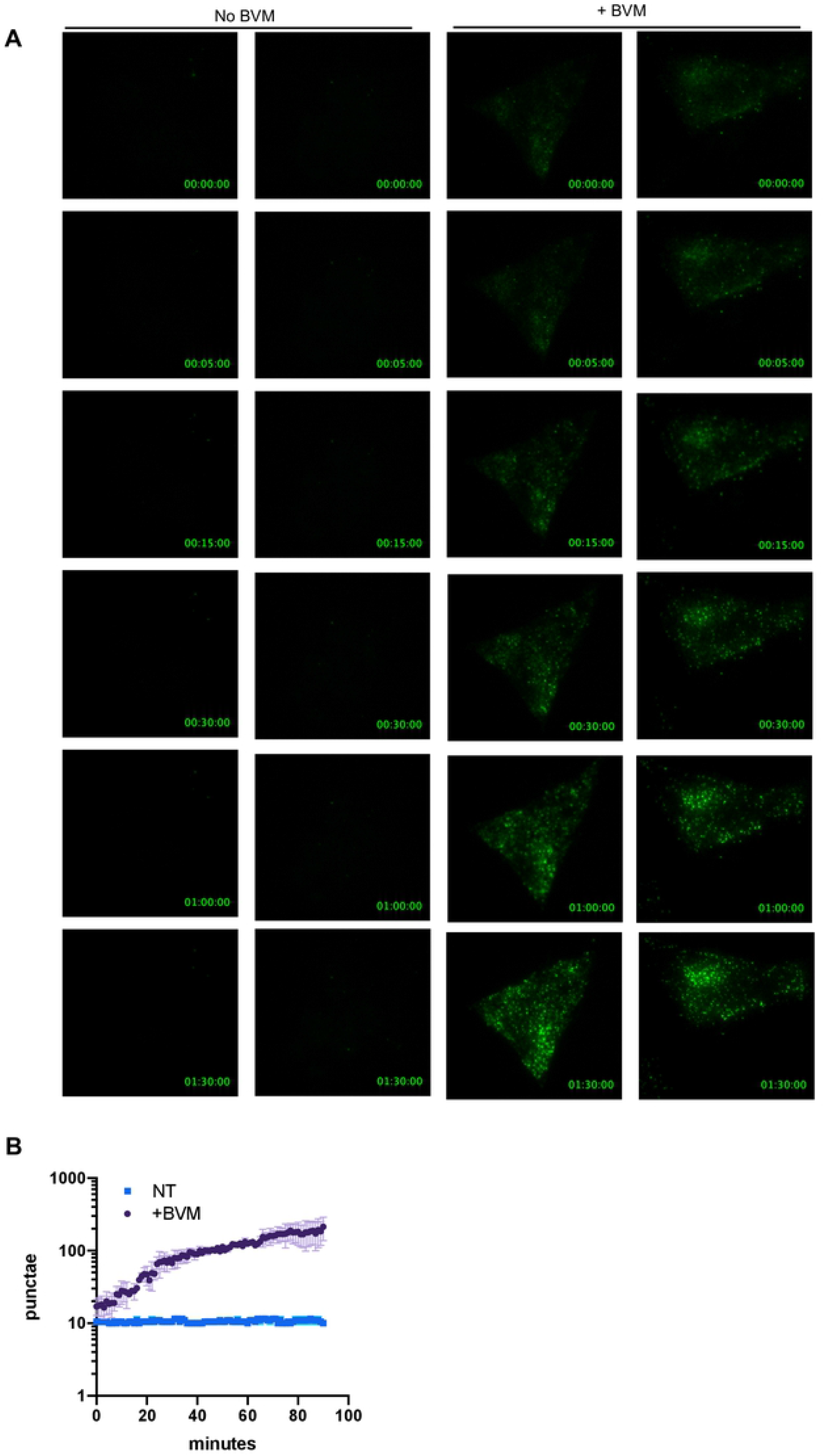
Visualization of Bevirimat-Induced assembly by TIRF-FM. (A) TIRF-FM microscopy of TZM cells infected with HIV-1_P289S_ NG. Representative time-lapse images of TZM cells treated with BVM at 28 hours post infection. Image labels: Hours:Minutes:Seconds post BVM addition. (B) Quantification of punctae per time point for images from (A).

## Discussion

Here, we report the identification of a single amino acid substitution (T371I) that rescues the replication of the defective, IP6-binding deficient mutant HIV-1_K359A_. Despite several attempts we were unable to generate revertant mutants for HIV-1_K290A_ and HIV-1_R18A_ (although follow-up studies demonstrated that the T371I substitution rescues HIV-1_K290A_ as well as HIV-1_K359A_). The inability to generate revertant mutants for HIV-1_R18A_ and HIV-1_K290A_ is likely due to more substantial impairment. Indeed, previous groups have shown that HIV-1_K290A_ is impaired to a greater extent than HIV-1_K359A_, potentially because K290 binds the 5 equatorial phosphates on IP_6_ while K359 coordinates the single axial phosphate, suggesting a greater role for K290 in coordinating IP_6_ (13). The inability to generate a revertant mutant rescuing HIV-1_R18A_ could be explained by functions of this residue in addition to coordinating IP_6_, such as recruitment the cellular protein FEZ1 or as service as a conduit for dNTPs into the mature core (10,11,20,21).

HIV-1_K359A/T371I_ is fully infectious despite containing a mutation (K359A) that renders VLPs unresponsive to IP_6_ *in vitro* and substantially impairs IP_6_ incorporation into virions (9,13). Indeed, we found no significant reduction in yield of HIV-1_K359A/T371I_ from IPMK KO 293T cells, in contrast to WT HIV-1. This finding suggests that HIV-1_K359A/T371I_ is no longer dependent on IP_6_ in virus producing cells, and that the lattice stabilizing mutation T371I can functionally substitute for IP_6_.

In addition to its role in promoting immature particle assembly, IP_6_ has also been implicated in stabilizing the mature lattice and promoting viral DNA synthesis by binding to a positively charged pore formed by R18 residues in the mature CA lattice (10,20). Previously it was proposed that the source of the IP_6_ required to stabilize the mature lattice in virions is selective recruitment by K290 and K359 residues, with IP_6_ being liberated to bind R18 residues following disruption of the immature lattice after proteolysis (9,10). The fact that HIV-1_K359A/T371I_ has no infectivity deficit, despite bearing a substitution known to decrease IP_6_ incorporation into virions argues against a requirement for IP_6_ post assembly. While we cannot formally exclude the possibility that HIV-1_K359A/T371I_ has regained the ability to efficiently package IP_6_, this is highly unlikely as 1) our data shows that HIV-1_K359A/T371I_ is not dependent on cellular IP_6_ levels in producer cells and 2) the T371I substitution does not result in a change to an amino acid with the appropriate properties (i.e., positively charged) to coordinate IP_6._ Furthermore, IPMK KO target MT4 cells were fully susceptible to infection by HIV-1_WT_, in agreement with previous studies (13). However, we also found no difference in the infectiousness of HIV-1_K359A/T371I_ in WT and IPMK KO MT4 target cells. If K359-dependent virion incorporation or target cells represent a source of IP_6_ required for post assembly functions, then impaired infectivity of HIV-1_K359A/T371I_ should be evident in IPMK KO target cells. However, our findings do not support the notion that IP_6_ is strictly required post assembly. This may reflect the possibility that other polyanions can fulfil a post assembly role. Indeed, recent studies demonstrated that other polyanions in mammalian cells such as glucose-1,6-bisphosphate can stabilize mature HIV-1 cores *in vitro* (22).

The T371I substitution, identified herein as a compensatory mutation that rescues infectivity deficit found in HIV-1_K359A_, has previously been reported to rescue the infectivity of virions containing substitutions that confer resistance to maturation inhibitors (MIs). These substitutions render HIV-1 assembly-defective in the absence of MIs (17) and the T371I substitution was shown to stabilize the CA-SP1 lattice in this context, effectively mimicking the action of MIs (16). Because the T371I substitution, apparently mimics the effect of MIs, we hypothesized that MIs might also rescue the infectivity of HIV-1_K359A_. Strikingly, we found that this was the case, and found that BVM can stimulate the assembly and release of HIV-1_K359A_. That the stabilizing effects of both the T371I substitution and BVM can compensate for the lack of IP_6_ coordination in HIV-1_K359A_ provides *in vivo* mechanistic support for the model proposed by Dick et al: i.e. that binding of IP_6_ to K290 and K359 residues stabilizes the immature lattice to drive particle assembly.

The ability to promote assembly via addition of a small molecule has potential utility in imaging and other studies of HIV-1 assembly. Indeed, when performing live cell time-lapse imaging, we observed BVM-induced assembly on the timescale of minutes. Such an experimental approach has potential utility for studying the sequence of events in HIV-1 viron assembly, such are RNA packaging, ESCRT protein recruitment, and protease activation, where virion assembly can be rapidly and (potentially) reversibly induced in real time simply by addition of a small molecule.

Together, these data support a model whereby stability of the immature CA lattice is finely tuned, with IP_6_ coordinating and stabilizing the otherwise repulsive positive charges of K290 and K359 to drive assembly. Manipulations that cause IP_6_ binding deficiency, either mutagenesis of K359 or decreasing IP_6_ levels in producer cells, destabilize the immature lattice and decrease production of progeny virions. Conversely, manipulations such as the T371I substitution or treatment of HIV-1_WT_ with BVM, hyper-stabilize the immature lattice in the wild type context and decrease HIV-1 infectivity. However, either the T371I substitution or BVM treatment are able to functionally substitute for IP_6_ in the context of HIV-1_K359A_ and rescue virion assembly and infectiousness. Understanding how small molecules such as BVM or IP_6_ can enhancing Gag lattice stability can provide new tools to study virion assembly and potential avenues for antiretroviral therapeutics.

## Materials and Methods

### Cells and media

293T cells were maintained in Dulbecco’s modified Eagle’s medium (DMEM, Gibco) supplemented with 10% fetal calf serum and gentamycin. MT4 cells were maintained in Roswell Park Memorial Institute (RPMI) 1640 Medium (Gibco) supplemented with 10% fetal calf serum and gentamycin. Cells were maintained at 37°C and 5% CO_2_. All transfections with viral plasmids were performed with polyethyleneimine.

### Plasmid construction

All full-length proviral plasmids used in this study were based on the HIV-1 clone NHG, a previously described HIV-1 clone that encodes GFP in place of Nef (Accession number: <u>JQ585717</u>)(23). Mutant viruses were derived from this parental plasmid using primer mutagenesis with fragments assembled into NHG digested with SpeI and SbfI using NEB HiFi DNA Assembly Master Mix according to the manufacturer’s instructions. Primers used for mutagenesis include: K359A F: 5’-CCGGCCATGCTGCAAGAGTTTTG; K359A R: 5’-CAAAACTCTTGCAGCATGGCCGG; T371I F: 5’-GCAATGAGCCAAGTAATAAATCCAGCTACC; T371I R: 5’-GGTAGCTGGATTTATTACTTGGCTCATTGC; P289S F: 5’-CATAAGACAAGGAAGTAAGGAACCCTTTAGAG; P289S R: 5’-CTCTAAAGGGTTCCTTACTTCCTTGTCTTATG; R57G F: 5’-TCAAAGTAGGACAGTATGATC; R57G R: 5’-GATCATACTGTCCTACTTTGA. The imaging construct HIV-1 NG was derived from NL4-3. The region of NL4-3 encoding Pol was deleted from bp 2294-4813 and a unique XbaI was added at bp 2301; Vpu was deleted and Env was truncated by removing bp 6056 – 7250 inserting NheI at 6056. These deletions and restriction sites were created through overlap PCR and cloned into NL4-3 via SphI and NheI (NL4-3 BssHII 5’-GCTGAAGCGCGCACGGCAAGAGGCG 5’-CTGAAGCGCGCACGGCAAGAGGCGAGG and dPol Xba AS 5’-CTACTATTCTTTCCCCTGCACTCTAGACTACTACTTTATTGTGACGAGGGGTCGC; dPol Xba S 5’-GCGACCCCTCGTCACAATAAAGTAGTAGTCTAGAGTGCAGGGGAAAGAATAGTAG and dVpudEnv NheI AS 5’-CTCCTCGCTAGCGTACTACTTACTGCTTTGATAG). The sequence encoding neon green (NG) was codon optimized to have nucleotide composition and codon usage similar to that of Pol using the Codon Optimization On-Line Tool from Singapore University (http://cool.syncti/org) and was synthesized by GeneArt. Neon Green was fused into the p6* frame of Pol through overlap PCR and inserted via SphI and XbaI (NL4-3 SphI 5’ AGTGCATGCAGGGCCTATTGCACC, Pol-NG AS 5’-CATGTTATCCTCCTCGCCCTTGCTCACCATCTTTATTGTGACGAGGGGTCGCTGCCA; Pol-NG S 5’-TGGCAGCGACCCCTCGTCACAATAAAGATGGTGAGCAAGGGCGAGGAGGATAACATG, NG XbaI 5’-CTCCTCTCTAGACTACTTGTACAGCTCGTCCATGCCCAT).

Mutagenesis of this construct was accomplished using the same primers as above.

### Viral stock production

293T cells were seeded at 6 × 10^6^ cells per 10cm dish and transfected the next day using polyethylenimine. 8 hours post transfection, cells were placed in fresh medium. For generation of full-length virus, 293T cells were transfected with 15 μg of proviral plasmid. For generation of imaging constructs, 293T cells were transfected with 6μg of proviral plasmid, 6μg of SYN-GP, and 1.2 μg of VSV-G. At 48 hours post transfection, supernatants were harvest and passed through a .22μM filter. Titer of full-length infectious viruses was determined by serial dilution on MT4 cells. At 48 hours post infection, cells were fixed with 4% PFA and assessed via flow cytometry. Titer of imaging constructs was determined by serial dilution on TZM-bl cells. 48 hours post infection, cells were fixed with 0.5% glutaraldehyde and stained with X-gal to visualize number of infected foci.

### Generation of IPMK KO cell lines

The IPMK-targeting guides g1: ATGTACGGGAAGGACAAAGT; g2: GGTGGACTCGATCGCCGGTG; or g3: CCGGCCACCTGATGCGAGAG were designed using the Broad Institute GPP Web Portal and cloned into lentiCRISPRv2 bearing a Hygromycin resistance cassette digested with BsmBI. lentiCRISPR v2 was a gift from Feng Zhang (Addgene plasmid # 52961; http://n2t.net/addgene:52961; RRID:Addgene_52961). VLPs were prepared as above, with the exception that 1 × 10^6^ 293Ts/well were seeded in a 6 well plate and transfected the next day with 1µg lentiCRISPRv2, 1µg of Gag-Pol, and 0.2 μg of VSV-G. At 48 hours post transduction of target cells with lentiCRISPRv2, cells were placed in selection with 100 μg/mL Hygromycin for ∼10-14 days. Single cell clones were obtained by limiting dilution, and editing was verified by amplifying and sequencing target loci using primers: IPMK Seq F: 5’-CGCTTCTGCTCTCCGTTATG and IPMK Seq R: 5’-GGATTTGGCGTGCACACCAG and assessment using Synthego ICE, which identifies Indel frequency in Sanger sequencing data (Synthego Performance Analysis, ICE Analysis. 2019. v2.0.). Control cells were obtained similarly, using a lentiCRISPRv2 plasmid not harboring a sgRNA cassette.

### Single cycle infectivity assays

WT control or IPMK KO 293T cells were seeded at 2.5 × 10^5^ cells/well in a 24 well plate and transfected with 625 ng of HIV-1_WT,_ HIV-1_K359A_, or HIV-1_K359A/T371I_ proviral plamids. Virions were prepared as above and titrated on MT4 cells. 24 hours post infection, Dextran Sulfate was added (50µg/mL) to limit infection to a single round. 48 hours post infection, cells were fixed with 4% PFA and assessed via flow cytometry.

### Western blotting

293Ts were seeded 5 × 10^5^ cells/well in a 12 well dish and transfected the next day with 1.25µg proviral plasmid. 48 hours post transfection, cell lysates and virions pelleted through 20% sucrose (14,000xg for 90 minutes at 4°C) were separated on a NuPage 4-12% Bis-Tris Gel (Invitrogen) and subsequently blotted onto a nitrocellulose membrane. Blots were blocked with Intercept Blocking Buffer (Li-Cor) and probed with primary antibody along with a corresponding IRDye 800CW- or IRDye 680-conjugated secondary antibody. Images were acquired using an Odyssey scanner (Li-Cor Biosciences). HIV-1 CA was detected using a human monoclonal anti-p24 (NIH AIDS Reagent Catalog #530).

### Imaging

5×10^4^ TZM-bl cells per well were plated in a Lab-Tek Chamber Slide and infected the following day with indicated imaging construct at an MOI of 1. For fixed samples, cells infected in the presence or absence of 5µM BVM were fixed 48 hours post infection and imaged on a DeltaVision OMX SR imaging system using a 60X Widefield oil immersion objective (Olympus) with an exposure time of 50ms, 10% Transmission, A488 nm laser. For live-cell samples, image acquisition began 26 hours post infection, with cells placed in the presence or absence of 5µM BVM at the time of image acquisition. Images were acquired at 37°C, 5% CO_2_ at indicated timepoints using a 60X Widefield oil immersion objective with an exposure time of 45ms, 5% Transmission A488 nm laser. For TIRF-FM, cells were imaged approximately 28-30 hours post infection in the presence or absence of 5µM BVM at 37°C, 5% CO_2_. Images were acquired every 1 min for 90 min using a 60X RING-TIRF-FM objective (Olympus Apo N 60X 1.49 Oil) with an exposure time of 100 ms, 10% Transmission A488 nm laser. Representative images were acquired, and all images were analyzed with FiJi (https://fiji.sc/)

### Graphing and statistical analysis

All graphs and corresponding statistical analyses were produced and analyzed with Graphpad Prism.

## Acknowledgments

We thank members of the Bieniasz lab for helpful discussions. This work was supported by grants from the NIAID; R01AI050111 and R01AI150998 (to PDB). DP was supported by a Medical Scientist Training Program grant from the NIGMS (T32GM007739) to the Weill Cornell/Rockefeller/Sloan Kettering Tri-Institutional MD-PhD Program and by the NIAID (F30AI157898). The content is solely the responsibilities of the authors and does not necessarily represent the official views of the National Institutes of Health.

## Supporting information

**S1-4 Movie: HIV-1 NG**_**P289S**_ **live cell widefield microscopy**

TZM cells infected with HIV-1 NG_P289S_ and imaged 26 hours post infection in the absence (Movie S1-2) or presence (Movie S3-4) of 5 µM BVM. Image acquisition began immediately after BVM addition. Images were acquired every 30 minutes. Movie labels: Hours:Minutes:Seconds post BVM addition.

**S5-6 Movie: HIV-1 NG**_**P289S**_ **live cell widefield microscopy at increased time resolution**

TZM cells infected with HIV-1 NG_P289S_ and imaged 26 hours post infection in the presence of 5 µM BVM. Image acquisition began immediately after BVM addition. Images were acquired every 3 minutes. Movie labels: Hours:Minutes:Seconds post BVM addition.

**S7-10 Movie: HIV-1 NG**_**P289S**_ **RING-TIRF-FM**

TZM cells infected with HIV-1_P289S_ NG and imaged 28 hours post infection in the presence (Movie S7-8) or absence (Movie S9-10) of 5µM BVM using RING-TIRM fluorescence microscopy. Image acquisition began immediately after BVM addition and images were acquired every 60 seconds. Image labels: Hours:Minutes:Seconds post BVM addition.

## References

1. Freed EO. HIV-1 assembly, release and maturation. Nat Rev Microbiol. 2015 Aug;13(8):484–96.

2. Dorfman T, Bukovsky A, Ohagen A, Höglund S, Göttlinger HG. Functional domains of the capsid protein of human immunodeficiency virus type 1. J Virol. 1994 Dec;68(12):8180–7.

3. Borsetti A, Ohagen A, Göttlinger HG. The C-terminal half of the human immunodeficiency virus type 1 Gag precursor is sufficient for efficient particle assembly. J Virol. 1998 Nov;72(11):9313–7.

4. Nermut MV, Hockley DJ, Jowett JB, Jones IM, Garreau M, Thomas D. Fullerene-like organization of HIV gag-protein shell in virus-like particles produced by recombinant baculovirus. Virology. 1994 Jan;198(1):288–96.

5. Jouvenet N, Neil SJD, Bess C, Johnson MC, Virgen CA, Simon SM, et al. Plasma membrane is the site of productive HIV-1 particle assembly. PLoS Biol. 2006 Dec;4(12):e435.

6. Coffin JM, Hughes SH, Varmus HE, editors. Retroviruses. Cold Spring Harbor (NY): Cold Spring Harbor Laboratory Press; 1997.

7. Sundquist WI, Kräusslich H-G. HIV-1 assembly, budding, and maturation. Cold Spring Harb Perspect Med. 2012 Jul;2(7):a006924.

8. Campbell S, Fisher RJ, Towler EM, Fox S, Issaq HJ, Wolfe T, et al. Modulation of HIV-like particle assembly in vitro by inositol phosphates. Proc Natl Acad Sci USA. 2001 Sep 11;98(19):10875–9.

9. Dick RA, Zadrozny KK, Xu C, Schur FKM, Lyddon TD, Ricana CL, et al. Inositol phosphates are assembly co-factors for HIV-1. Nature. 2018 Aug 1;560(7719):509–12.

10. Mallery DL, Márquez CL, McEwan WA, Dickson CF, Jacques DA, Anandapadamanaban M, et al. IP6 is an HIV pocket factor that prevents capsid collapse and promotes DNA synthesis. Elife. 2018 May 31;7.

11. Xu C, Fischer DK, Rankovic S, Li W, Dick R, Runge B, et al. Permeability of the HIV-1 capsid to metabolites modulates viral DNA synthesis. BioRxiv. 2020 May 1;

12. Yu A, Lee EMY, Jin J, Voth GA. Atomic-scale characterization of mature HIV-1 capsid stabilization by inositol hexakisphosphate (IP6). Sci Adv. 2020 Sep 16;6(38).

13. Mallery DL, Faysal KMR, Kleinpeter A, Wilson MSC, Vaysburd M, Fletcher AJ, et al. Cellular IP6 Levels Limit HIV Production while Viruses that Cannot Efficiently Package IP6 Are Attenuated for Infection and Replication. Cell Rep. 2019 Dec 17;29(12):3983–3996.e4.

14. Ricana CL, Lyddon TD, Dick RA, Johnson MC. Primate lentiviruses require Inositol hexakisphosphate (IP6) or inositol pentakisphosphate (IP5) for the production of viral particles. PLoS Pathog. 2020 Aug 10;16(8):e1008646.

15. Dick RA, Xu C, Morado DR, Kravchuk V, Ricana CL, Lyddon TD, et al. Structures of immature EIAV Gag lattices reveal a conserved role for IP6 in lentivirus assembly. PLoS Pathog. 2020 Jan 27;16(1):e1008277.

16. Fontana J, Keller PW, Urano E, Ablan SD, Steven AC, Freed EO. Identification of an HIV-1 Mutation in Spacer Peptide 1 That Stabilizes the Immature CA-SP1 Lattice. J Virol. 2016 Jan 15;90(2):972–8.

17. Waki K, Durell SR, Soheilian F, Nagashima K, Butler SL, Freed EO. Structural and functional insights into the HIV-1 maturation inhibitor binding pocket. PLoS Pathog. 2012 Nov 8;8(11):e1002997.

18. Keller PW, Adamson CS, Heymann JB, Freed EO, Steven AC. HIV-1 maturation inhibitor bevirimat stabilizes the immature Gag lattice. J Virol. 2011 Feb;85(4):1420–8.

19. Nguyen AT, Feasley CL, Jackson KW, Nitz TJ, Salzwedel K, Air GM, et al. The prototype HIV-1 maturation inhibitor, bevirimat, binds to the CA-SP1 cleavage site in immature Gag particles. Retrovirology. 2011 Dec 7;8:101.

20. Jacques DA, McEwan WA, Hilditch L, Price AJ, Towers GJ, James LC. HIV-1 uses dynamic capsid pores to import nucleotides and fuel encapsidated DNA synthesis. Nature. 2016 Aug 18;536(7616):349–53.

21. Huang P-T, Summers BJ, Xu C, Perilla JR, Malikov V, Naghavi MH, et al. FEZ1 Is Recruited to a Conserved Cofactor Site on Capsid to Promote HIV-1 Trafficking. Cell Rep. 2019 Aug 27;28(9):2373–2385.e7.

22. Dostálková A, Kaufman F, Křížová I, Vokatá B, Ruml T, Rumlová M. In vitro quantification of the effects of IP6 and other small polyanions on immature HIV-1 particle assembly and core stability. J Virol. 2020 Jul 29;

23. Wilson SJ, Schoggins JW, Zang T, Kutluay SB, Jouvenet N, Alim MA, et al. Inhibition of HIV-1 particle assembly by 2’,3’-cyclic-nucleotide 3’-phosphodiesterase. Cell Host Microbe. 2012 Oct 18;12(4):585–97.

